# Regional Brain Entropy, Brain Network and Structural-Functional Coupling in Human Brain

**DOI:** 10.1101/2024.11.14.623700

**Authors:** Donghui Song

**Affiliations:** State Key Laboratory of Cognitive Neuroscience and Learning, Beijing Normal University, Beijing,100091, China; IDG/McGovern Institute for Brain Research, Beijing Normal University, Beijing,100091, China

**Keywords:** Brain entropy (BEN), Brain network, Resting-state fMRI, Diffusion tensor imaging (DTI), Gradients

## Abstract

Brain entropy (BEN) indicates the irregularity, unpredictability and complexity of brain activity. In healthy brains, resting-state fMRI-based regional BEN (rBEN) distribution has been shown to have potential relationships with functional brain networks. However, the relationship between rBEN and structural and functional networks, as well as how rBEN facilitates the coupling between structural and functional networks remains unclear. Additionally, network dimensionality reduction methods, such as the use of connectome gradients, have become popular in recent years for explaining the hierarchical architecture of brain function, and the sensorimotor-association (S-A) axis, as a major axis of hierarchical cortical organization, exerts a widespread influence on both brain structure and function. However, how this functional hierarchy affects the relationship between rBEN and brain networks remains unclear. In this study, we systematically examine the relationship between rBEN and both structural and functional networks, including BOLD-based functional networks and MEG-based functional networks. We also assess the impact of the BEN and network gradients, as well as the influence of the S-A axis, on the relationship between rBEN and brain networks.

Our results reveal a negative correlation between BEN and the network efficiency of both structural and BOLD-based functional networks, as assessed by average connection strength, degree centrality, and local efficiency. Additionally, BEN shows a negative correlation with the coupling between structural and BOLD functional networks. In contrast, the relationship between BEN and MEG functional network efficiency varies from negative to positive across different frequency bands, with a shift from negative to positive correlation observed in the coupling between BEN and MEG-functional networks. Moreover, the coupling between BOLD and MEG functional networks is positively correlated with BEN. The relationship between BEN and network gradients is complex, and no consistent patterns were observed. Importantly, the relationship between BEN and brain networks is influenced by the S-A axis, and the connection between BEN and networks is further modulated by changes in cytoarchitectural organization.

These results are consistent with the commonly observed relationship between elevated rBEN and impaired networks in psychiatric disorders. The negative correlation between BEN and the efficiency of both structural and BOLD functional networks, as well as their coupling, may suggest that lower rBEN is associated with higher information processing potential. Additionally, the positive correlation between BEN and high-frequency MEG functional networks, as well as the coupling between BOLD and MEG functional networks, may indicate that brain processes related to information acquisition and integration could increase rBEN. In summary, this complex relationship may reflect a dynamic process in which the brain continuously acquires external information for integration (increasing entropy) and internalizes it (decreasing entropy) as part of its information-processing mechanism in different timescales.

## 1. Introduction

It is well known that the human brain consists of about 100 billion interconnected neurons (Herculano-Houzel 2009, Von Bartheld, Bahney et al. 2016). The collective activity of these neurons forms regional neural circuits, which in turn contribute to larger-scale brain networks (Bassett and Sporns 2017, Schröter, Paulsen et al. 2017, Luo 2021). Current research on the human brain has established a connectomics framework that spans microscopic, mesoscopic, and macroscopic levels (Bassett and Sporns 2017). However, an important yet often overlooked question is how large-scale brain networks emerge from regional neural activity, and the connection between regional neural activity and brain networks has yet to be established well. Moreover, the coupling between structure and function has long been an important issue in neuroscience, whether regional neural activity facilitates the coupling between these structural and functional networks remains unclear. In this study, we utilize regional brain entropy (rBEN) (Wang, Li et al. 2014), brain networks (Bellec, Perlbarg et al. 2006, Brookes, Hale et al. 2011, Power, Cohen et al. 2011, Park and Friston 2013), and connectivity gradients (Margulies, Ghosh et al. 2016) to build a bridge between regional neural activity and brain networks and assess whether regional neural activity promotes coupling between structural and functional networks.

Entropy originated in thermodynamics, where Clausius used it to describe the second law of thermodynamics, stating that the entropy will always increase or stay the same over time in an isolated system(Clausius 1857). Later, Shannon introduced the concept of entropy in information theory, as a measure of the average amount of information in a message or the uncertainty about the source of information (Shannon 1948). This concept has gradually evolved into a powerful metric for measuring physiological signals (Pincus 1991, Richman and Moorman 2000, Lake, Richman et al. 2002, Pincus 2006, Sabeti, Katebi et al. 2009, Gómez and Hornero 2010, Yao, Lu et al. 2013, Wang, Li et al. 2014, Chang, Song et al. 2018, Lin, Lee et al. 2019, Song, Chang et al. 2019, Song, Chang et al. 2019, Xue, Yu et al. 2019, Liu, Song et al. 2020, Wang and Initiative 2020, Wang 2021, Jiang, Cai et al. 2023, Song Donghui 2023, Del Mauro and Wang 2024, Dong-Hui Song 2024, Song, Deng et al. 2024, Song and Wang 2024, Song and Wang 2024, Zhao, Shuai et al. 2024) . In the human brain, electrophysiological signals, including electroencephalography (EEG) and magnetoencephalography (MEG), have been extensively used to assess brain entropy (Stam 2005, Pincus 2006, Lau, Pham et al. 2022), which has demonstrated the ability to distinguish among various neurological and psychiatric disorders (Sabeti, Katebi et al. 2009, Gómez and Hornero 2010, Faust, Ang et al. 2014, Lau, Pham et al. 2022). However, its effectiveness is often limited due to low spatial resolution and high noise, which can obscure regional neural activity information. In contrast, functional magnetic resonance imaging (fMRI) offers excellent spatial resolution, allowing us to gain deeper insights into regional neural activity information through hemodynamics. About ten years ago, Ze Wang et al (2014) proposed a method for measuring rBEN using sample entropy (Richman and Moorman 2000, Lake, Richman et al. 2002) based on fMRI and mapped the hierarchical distribution of rBEN in healthy brains (Wang, Li et al. 2014) . The rBEN has now evolved into a relatively mature metric for measuring regional brain activity. The rBEN not only reflects information about regional spontaneous brain activity (Song, Chang et al. 2019) but is also related to brain structure (Del Mauro and Wang 2024, Song and Wang 2024) and neurochemical architecture (Song and Wang 2024, Song and Wang 2024). Furthermore, it can predict individual cognitive performance (Wang 2021, Lin, Chang et al. 2022, Camargo, Del Mauro et al. 2024, Del Mauro and Wang 2024, Song and Wang 2024) and behavioral characteristics (Song and Wang 2024) and is highly sensitive to neural modulation (Chang, Zhang et al. 2018, Song, Chang et al. 2019, Liu, Song et al. 2024, Song, Deng et al. 2024). Additionally, it is associated with changes in brain activity related to various neuropsychiatric disorders and their treatments (Chang, Song et al. 2018, Chang, Zhang et al. 2018, Lin, Lee et al. 2019, Xue, Yu et al. 2019, Liu, Song et al. 2020, Wang and Initiative 2020, Song Donghui 2023, Dong-Hui Song 2024).

Modern neuroimaging techniques make available to reconstruct structural connectivity (SC) networks by tracing tractography using diffusion MRI (Hagmann, Kurant et al. 2007, Yeh, Jones et al. 2021), thereby representing the physical connections between neural elements. SC promotes electrical and chemical signals to establish connections between distant neural populations, thereby forming functional connectivity (FC) networks (Suárez, Markello et al. 2020). The most widely used method for reconstructing FC networks is based on time series synchronicity from resting-state fMRI. FC networks reconstructed from resting-state fMRI exhibit a moderate coupling with structural networks at whole brain, with spatial heterogeneity in this coupling. Specifically, there is a higher coupling in unimodal cortex and a lower coupling in multimodal cortex (Suárez, Markello et al. 2020, Shafiei, Baillet et al. 2022, Hansen, Shafiei et al. 2023, Liu, Shafiei et al. 2023). This pattern is generally thought to facilitate functional flexibility. Similarly, FC can also be reconstructed from electrophysiological data, including EEG and MEG. However, recent studies have found that FC derived from MEG does not align spatially with BOLD-driven FC. The correspondence is stronger in unimodal cortex but weaker in multimodal cortex. Furthermore, BOLD-driven FC cannot be fully explained by a single frequency band of MEG-derived FC, with the beta band contributing the most (Shafiei, Baillet et al. 2022, Hansen, Shafiei et al. 2023, Liu, Shafiei et al. 2023).

Although the structural networks, functional networks, and their coupling relationships have been widely studied, the relationship between these networks and rBEN remains unclear. Our previous research has shown that rBEN exhibits more consistent variations within large-scale brain networks than across brain regions, suggesting a potential link between rBEN and FC networks (Song and Wang 2024). In the resting state, lower resting state rBEN (rsBEN) is associated with higher intelligence and better educational attainment in the default mode network (DMN) and executive control network (ECN), which are typically linked to stronger functional connectivity (Song, Zhou et al. 2008, Hearne, Mattingley et al. 2016). Therefore, it can be hypothesized that rBEN may be negatively correlated with FC strength and lower rsBEN might be associated with enhanced information processing potential capacity (Wang 2021, Wang 2021, Lin, Chang et al. 2022). In addition to directly analyzing brain networks, recent conceptual and methodological developments allow macroscale brain networks to be mapped to low dimensional manifold representations, also described as gradients (Margulies, Ghosh et al. 2016, Vos de Wael, Benkarim et al. 2020). Gradient analysis of connectivity data has been extensively applied to structural and functional networks from MRI in specific brain regions, including the neocortex (Margulies, Ghosh et al. 2016, Vázquez-Rodríguez, Suárez et al. 2019, Wang 2020), subcortex (Tian, Margulies et al. 2020), and cerebellum (Guell, Schmahmann et al. 2018). And our recent research indicates that rBEN under movie-watching conditions (mvBEN) shows higher test-retest reliability (Song and Wang 2024). We also assess the relationship between rBEN under movie-watching conditions and SC, FC and connectivity gradients.

## 2. Methods

We have computed resting-state rBEN maps (rsBEN) and movie-watching rBEN maps (mvBEN) for 176 participants from our previous studies (Song and Wang 2024, Song and Wang 2024) using HCP 7T release (Van Essen, Smith et al. 2013). The whole brain rBEN maps were parcellated into 400 parcels based on the Schaefer 400 (Schaefer, Kong et al. 2018), and group average rBEN maps were computed from 176 participants (Song and Wang 2024). Additionally, we obtained the mean SC matrix, as well as the mean FC matrices based on both BOLD and MEG from 33 participants in the HCP Young Adult dataset at https://github.com/netneurolab/shafiei_megfmrimapping/tree/main/data (Shafiei, Baillet et al. 2022). For the data processing and construction methods of BOLD-FC and MEG-FC networks, please refer to the original article (Shafiei, Baillet et al. 2022, Liu, Shafiei et al. 2023).

## 3. Results

### 3.1 The relationship of rBEN and average connection strength

Each connectivity matrix is averaged by row to calculate the average connection strength for each node, ultimately forming the average structural connection strength, as well as the average BOLD and MEG functional connectivity strength for the 400 nodes. Then, the correlation between rBEN and the average connection strength is computed using Pearson’s correlation coefficients. The results show that neither rsBEN (r400=0.05, p>0.05) nor mvBEN (r400=-0.06, p>0.05) exhibit a significant correlation with the average structural connection strength. However, both rsBEN (r400=-0.25, p<0.001) and mvBEN (r400=-0.40, p<0.001) show a significant negative correlation with the average BOLD-FC strength. The rsBEN and mvBEN exhibited a negative correlation with the average MEG FC strength in the low-frequency bands ((delta (2 to 4 Hz), theta (5 to 7 Hz), alpha (8 to 12 Hz) and beta (15 to 29 Hz)), which gradually shifted to a positive correlation with the average MEG FC strength in the high-frequency bands,(low gamma (30 to 59 Hz), and high gamma (60 to 90 Hz)) (Fig 1).

**Fig 1.**
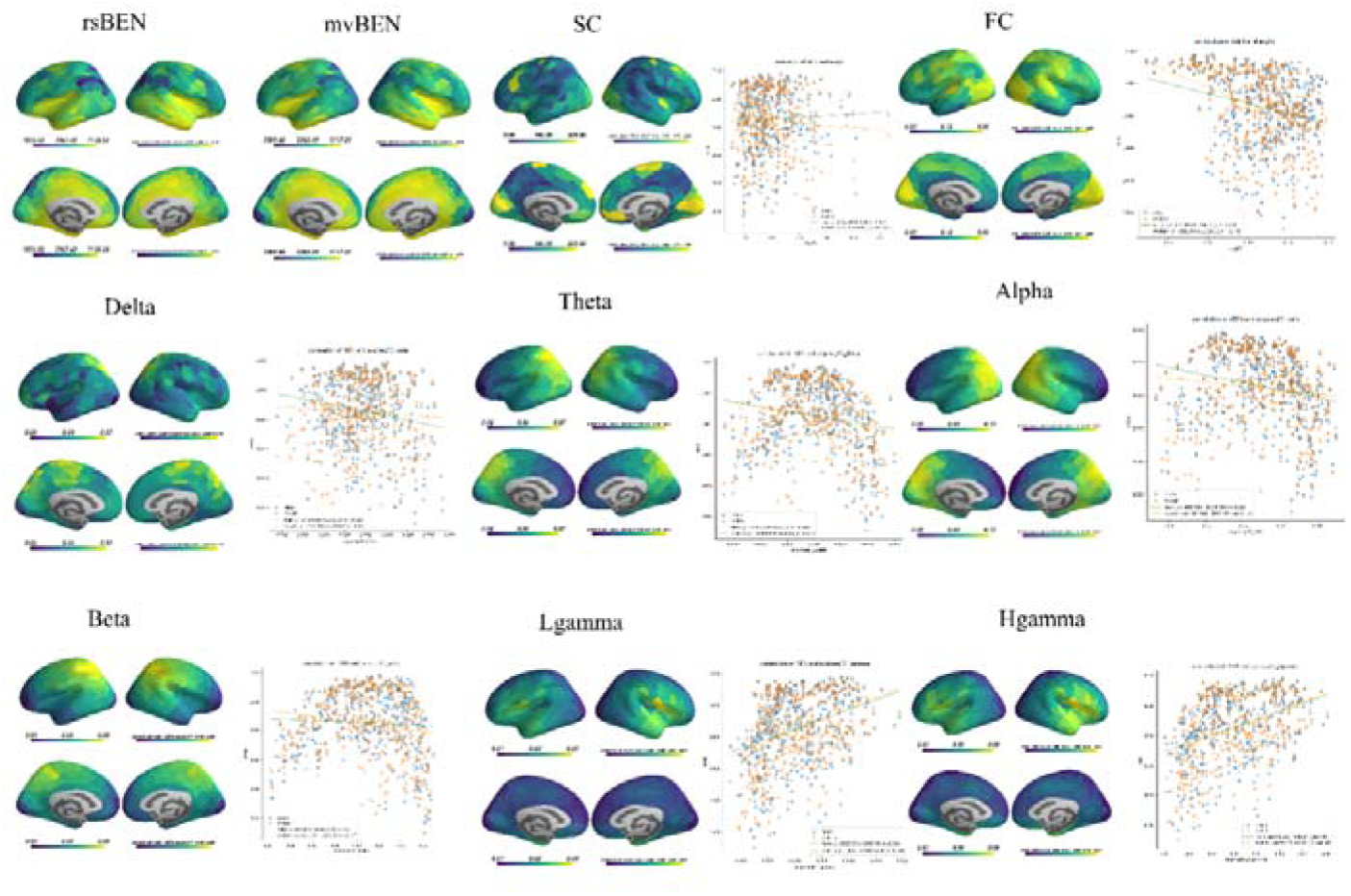
The relationship between rBEN and average connection strength. From top to bottom and left to right, the figures show the group-average rsBEN, mvBEN, the average connection strength of each node in SC, BOLD-FC, and MEG-FC across different frequency bands. The scatter plots display the spatial correlation between rsBEN, mvBEN, and the average connection strength of each network, with linear fitting applied.

### 3.2 The relationship of rBEN, degree centrality (DC) and local efficiency (LE)

To assess the relationship between rBEN and local brain network efficiency, we constructed both structural and functional networks, using degree centrality (DC) and local efficiency (LE) metrics from graph theory to characterize the local efficiency of the networks. We found that both rsBEN and mvBEN were significantly negatively correlated with the DC and LE of the structural network, as well as with the BOLD-based FC. Additionally, rsBEN and mvBEN showed a negative correlation with MEG-driven FC in low-frequency bands (delta (2 to 4 Hz), theta (5 to 7 Hz), which gradually shifted to a positive correlation in high-frequency bands (alpha (8 to 12 Hz) and beta (15 to 29 Hz), low gamma (30 to 59 Hz), and high gamma (60 to 90 Hz) (Fig 2).

**Fig 2.**
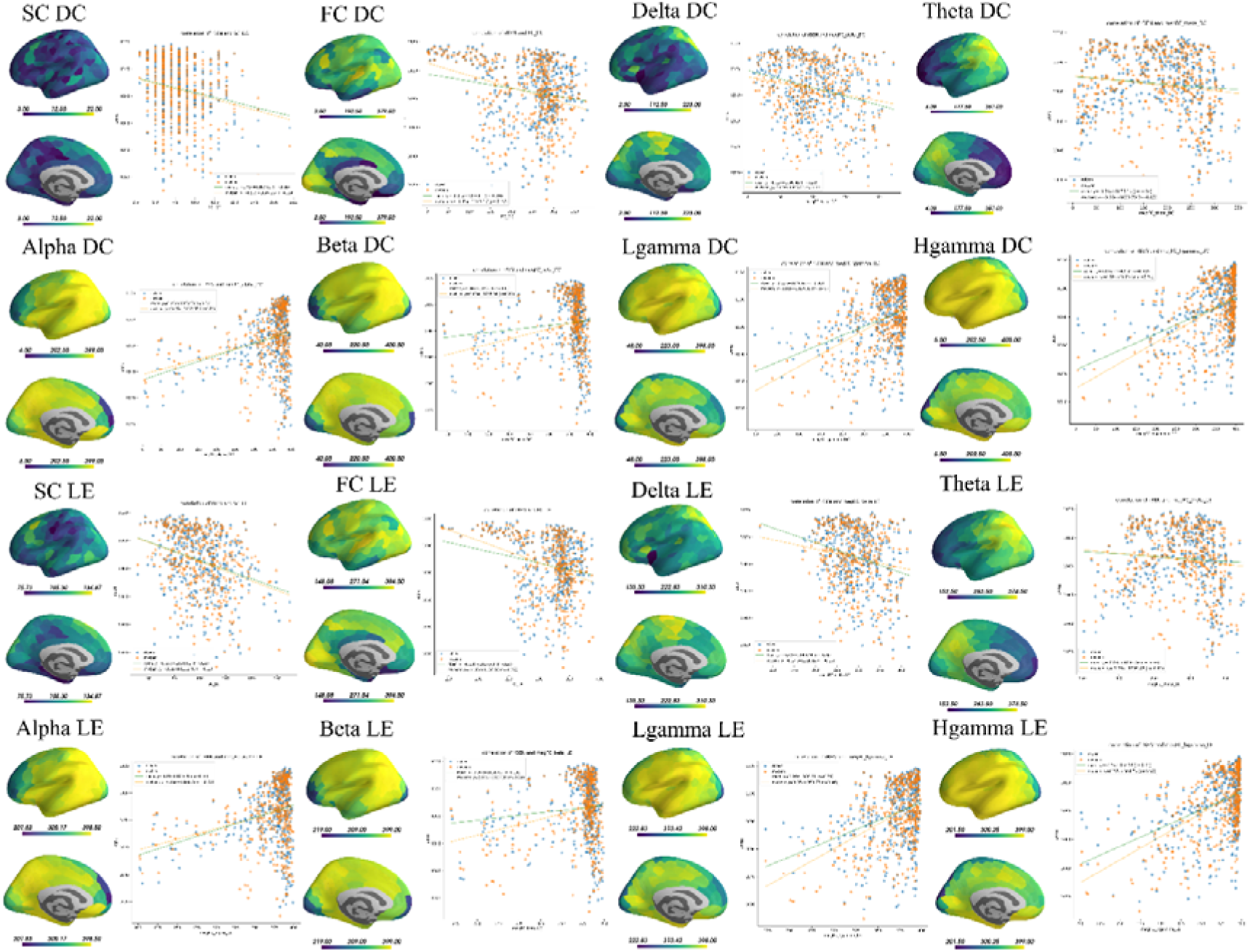
The relationship of rBEN, degree centrality (DC) and local efficiency (LE). From top to bottom and left to right, the figures show the DC and LE of each node in SC, BOLD-FC, and MEG-FC across different frequency bands. The scatter plots display the spatial correlation between rsBEN, mvBEN, and the DC and LE of each network, with linear fitting applied.

### 3.3 The relationship between rBEN and connectivity gradients

Gradient 1 of the structural connectivity, which spans from lateral to medial and from left to right, did not show any significant correlation with rsBEN or mvBEN. Gradient 2, which extends from anterior to posterior, displayed a negative correlation with mvBEN (r400 = -0.17, p < 0.001). For BOLD-FC, Gradient 1, which transitions from unimodal to multimodal regions, showed a negative correlation with rsBEN (r400 = -0.12, p < 0.05). Gradient 2, which spans from motor cortex to the association cortex and then to the visual cortex, showed a negative correlation with mvBEN (r400 = -0.38, p < 0.001). In contrast, the gradients of MEG-FC across different frequency bands varied, with the most prominent correlations observed with mvBEN (Fig 3).

**Fig 3.**
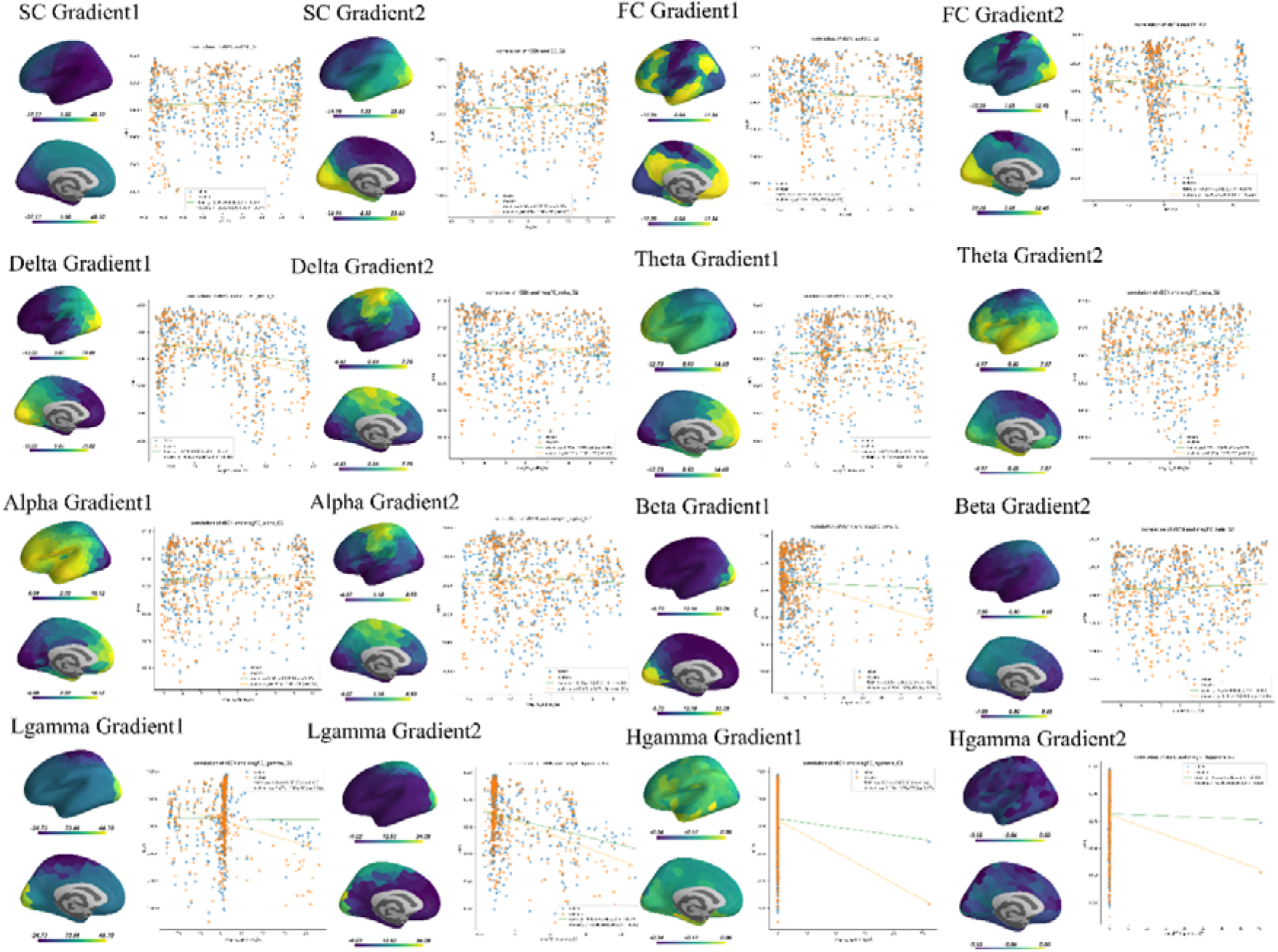
The relationship between rBEN and connectivity gradients. From top to bottom and left to right, the figures show the connectivity gradients of SC, BOLD-FC, and MEG-FC across different frequency bands. The scatter plots display the spatial correlation between rsBEN, mvBEN, and the DC and LE of each network, with linear fitting applied.

### 3.4 The relationship between rBEN, structural-functional coupling (SFC) and Bold-MEG coupling

We computed the local coupling between the structural and functional networks using correlation coefficients at each node. We found that both rsBEN (r400 = -0.47, p < 0.001) and mvBEN (r400 = -0.56, p < 0.001) were significantly negatively correlated with the coupling strength between the structural network and BOLD-FC. Additionally, rsBEN showed a significant negative correlation with the coupling between the structural network and MEG-FC in the low-frequency bands, which shifted to a significant positive correlation in the high-frequency bands. Both rsBEN and mvBEN exhibited significant positive correlations with the coupling between BOLD-FC and MEG-FC(Fig 4).

**Fig 4.**
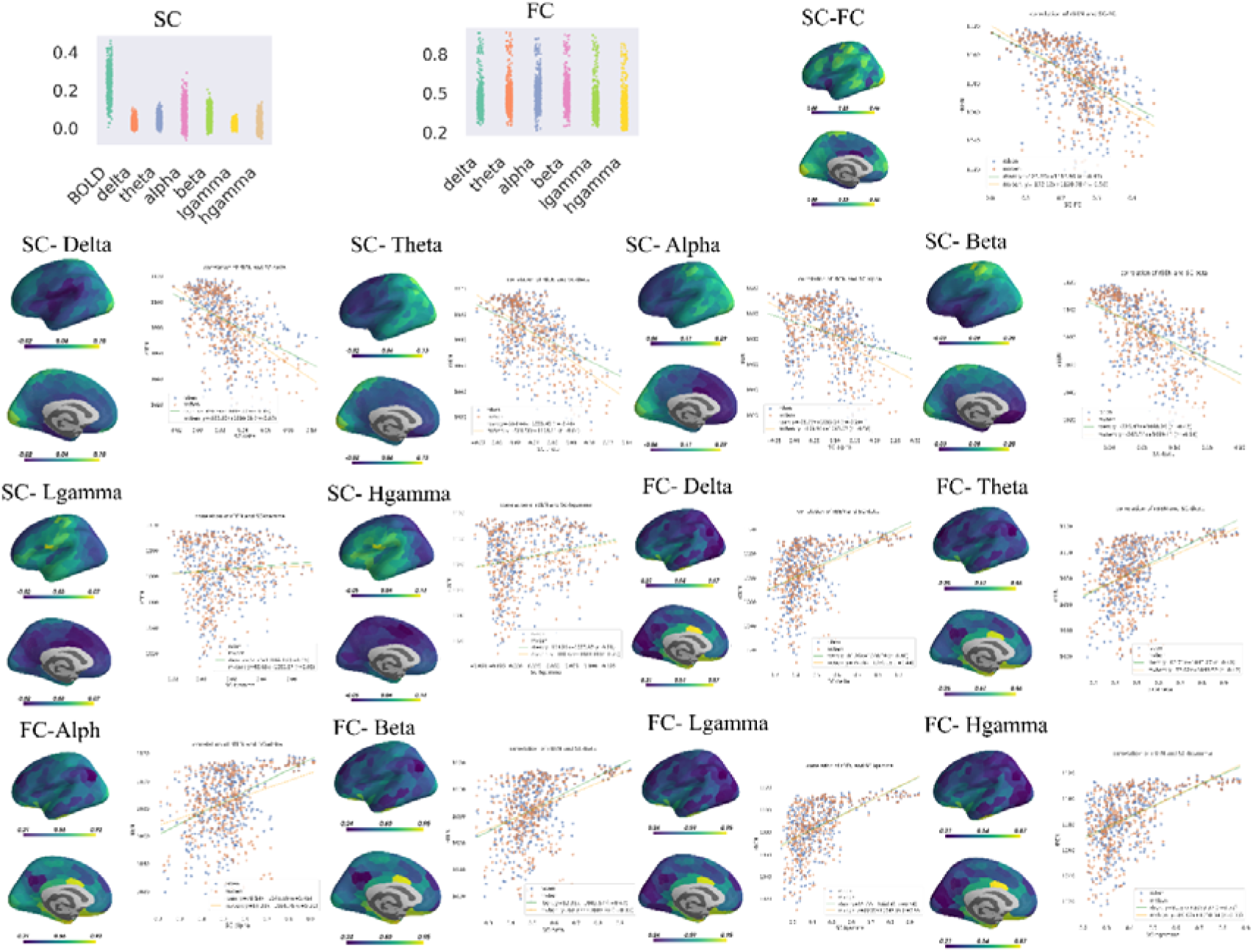
The relationship between rBEN and SFC, BOLD-FC-MEG-FC coupling. From top to bottom and left to right, the figures show the coupling strength between SC and BOLD-FC, and between SC and MEG-FC at each node, as well as the coupling strength between BOLD-FC and MEG-FC at each node. These coupling strengths are then mapped onto the brain atlas. The scatter plots display the spatial correlation between rsBEN, mvBEN, and coupling strength, with linear fitting applied.

### 3.5 The relationship between rBEN, average connection strength, SFC, Bold-MEG coupling and cytoarchitectural organization

We performed correlation analyses between rBEN, average connection strength, and structural-functional coupling with cell-type sampling across 50 layers. We found that rBEN, average connection strength, and structural-functional coupling were all associated with the cellular layer organization (Fig 5).

**Fig 5.**
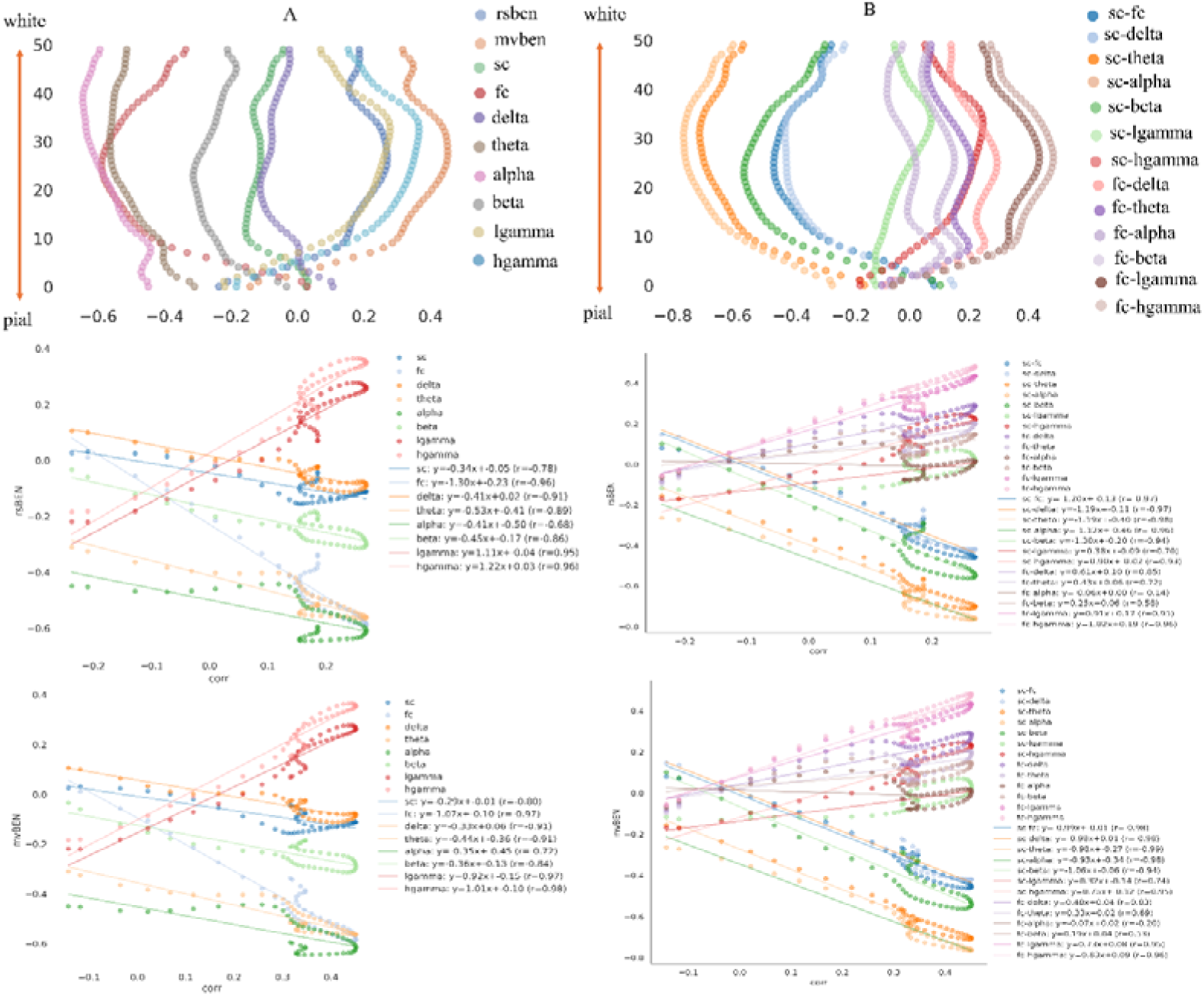
The relationship between rBEN, average connection strength, SFC, Bold-MEG coupling and cytoarchitectural organization.

**Fig 6.**
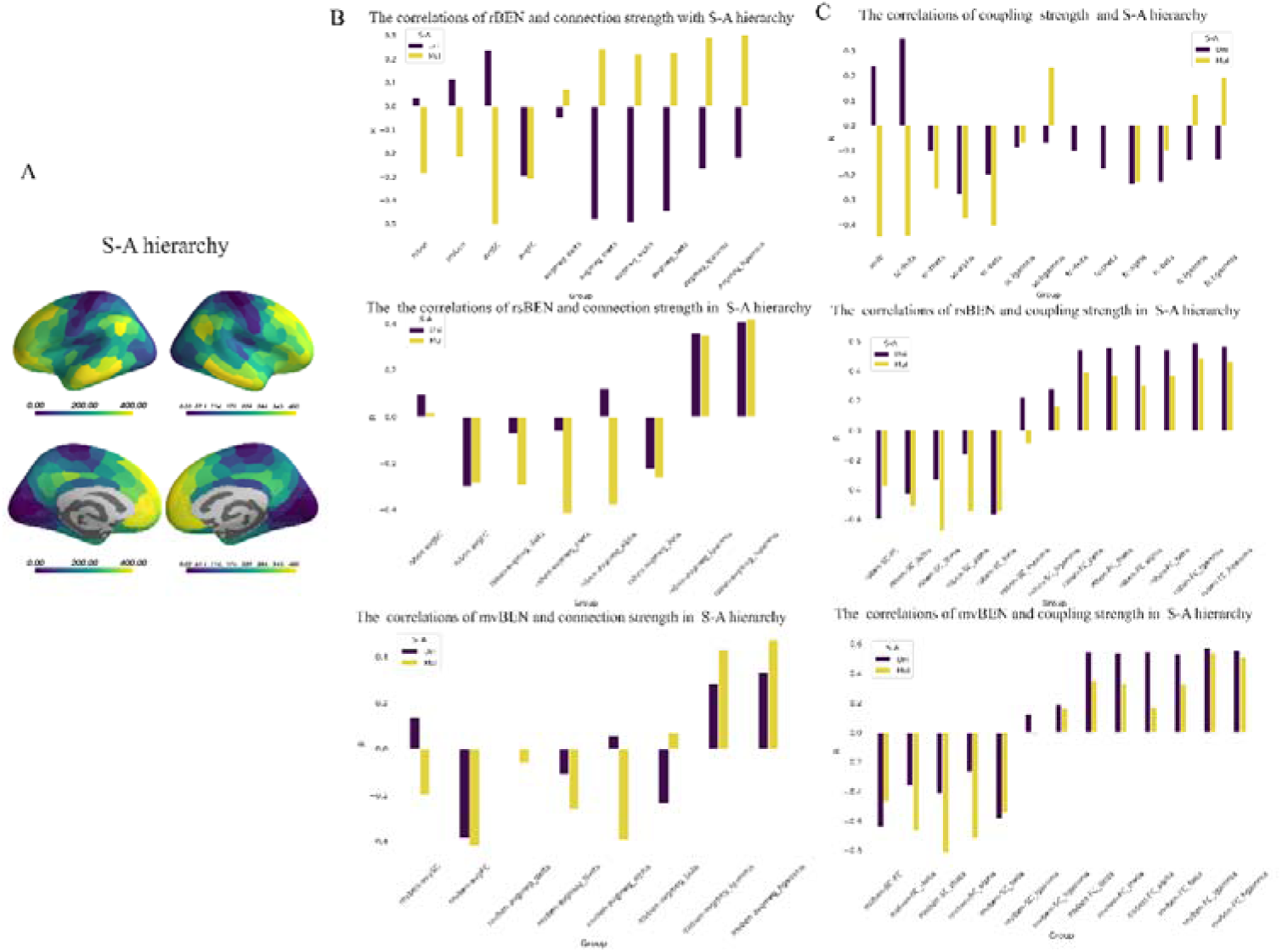
The relationship between rBEN, average connection strength and SFC long sensorimotor-association hierarchy

**Fig 7.**
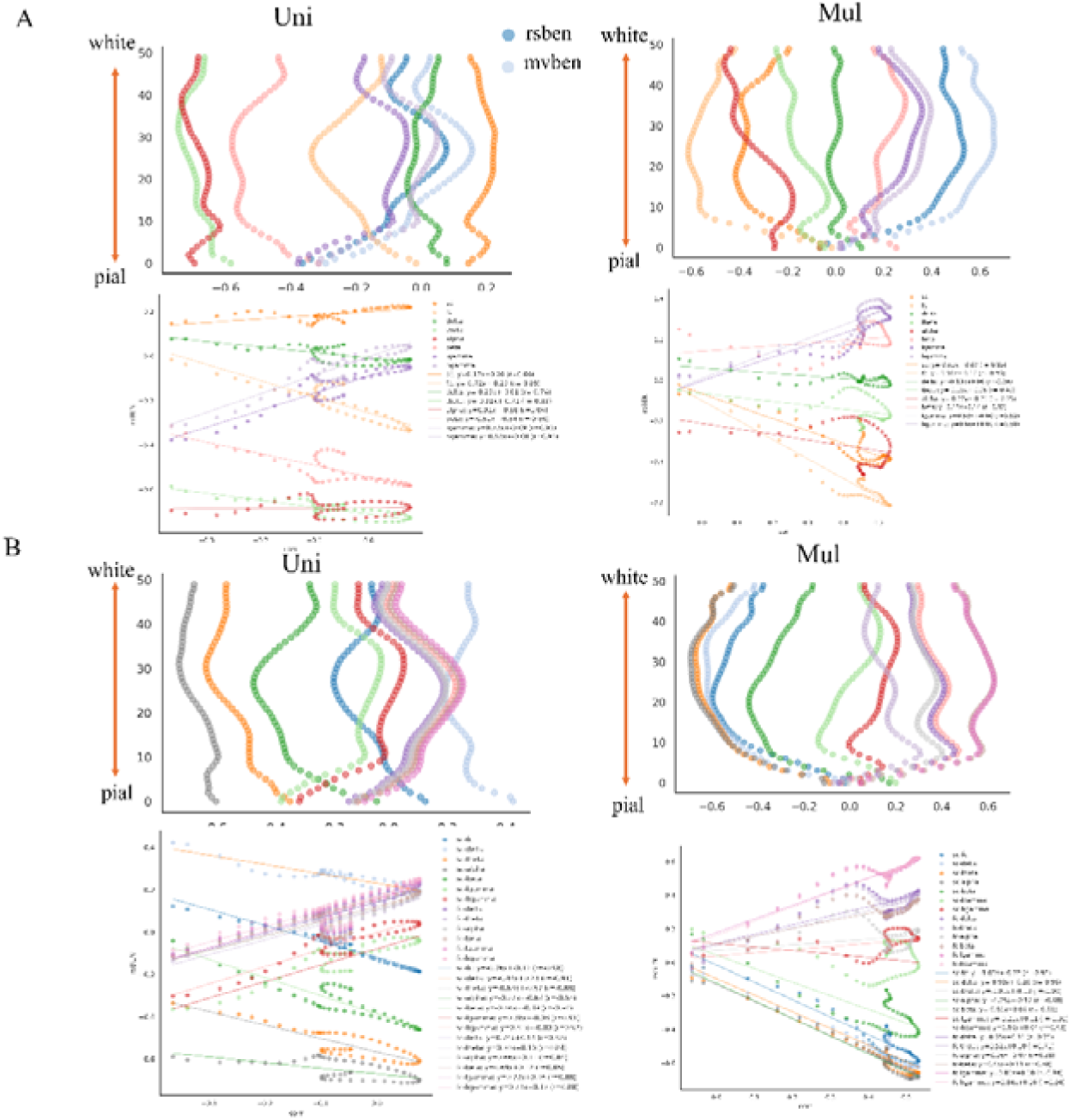
The relationship between rBEN, average connection strength, SFC, BOLD-MEG coupling and cytoarchitectural organization along sensorimotor-association hierarchy

### 3.6 The relationship between rBEN, average connection strength, SFC, Bold-MEG coupling and cytoarchitectural organization along sensorimotor-association hierarchy

We also performed an analysis of the sensorimotor-association (S-A) axis and found that rBEN, average connectivity strength of SC, as well as the average connectivity strength of MEG-FC, and the structural-functional coupling relationship, are all related to the hierarchical structure of the S-A axis. This, in turn, affects the correlations between rBEN and these measures. Furthermore, an analysis of the cytoarchitectural organization along the S-A axis revealed that the average connectivity strength of SC shows completely opposite correlation patterns with the cytoarchitectural organization in the unimodal cortex and multimodal cortex: it exhibits a positive correlation in the unimodal cortex and a negative correlation in the multimodal cortex.

## 4. Discussion

The local efficiency of SC and BOLD-FC with rBEN show a significant negative correlation, and rBEN also negatively correlates to coupling SC-BOLD-FC, suggesting that lower rsBEN may be associated with potential capacity in information processing in the human brain. This perspective is supported by our previous findings on brain entropy and related brain network studies. In our prior research, we demonstrated that lower rsBEN in the fronto-parietal region was negatively correlated with intelligence and education level (Wang 2021), while the network efficiency in this region showed a positive correlation with intelligence (Van Den Heuvel, Stam et al. 2009, Langer, Pedroni et al. 2012). Additionally, network efficiency is positive correlation with cognition and task performance (Achard and Bullmore 2007, Park and Friston 2013), rsBEN is negatively correlated with cognitive performance (Del Mauro and Wang 2024) and task-related activation and deactivation(Lin, Chang et al. 2022). In clinical, a large body of research has shown that disruptions in the brain networks are linked to various psychiatric disorders (Menon 2011, Ajilore, Lamar et al. 2014), while increased rBEN in various psychiatric disorders (Li, Fang et al. 2016, Lin, Lee et al. 2019, Xue, Yu et al. 2019, Liu, Song et al. 2020, Wang and Initiative 2020, Jiang, Cai et al. 2023, Dong-Hui Song 2024).

The reversal of the relationship between rBEN and different frequency bands of MEG, along with the positive correlation between rBEN and coupling of BOLD-FC and MEG-FC, may indicate that in the process of rapid information processing, the generation of entropy outweighs the process of reducing entropy. Our previous research has shown that rBEN increases in associate cortex and decreases in audiovisual cortex during movie-watching (Song and Wang 2024), and the pattern is inversely related to the inter-subject correlation in strength (Gruskin and Patel 2022), suggesting that faster processing of sensory information occurs in audiovisual cortex, while more refined and slower information processing in higher-order associative cortices.

This inverse relationship may suggest differences in the timescales of brain information processing (Golesorkhi, Gomez-Pilar et al. 2021, Chang, Nastase et al. 2022, Wolff, Berberian et al. 2022, Catal, Wolman et al. 2023, Senkowski and Engel 2024). We observed that rsBEN is negatively correlated along the multimodal cortex, while mvBEN, SC connection strength, and SC-BOLD FC coupling along the S-A axis exhibit an inverted correlation pattern. Specifically, they show a positive correlation in unimodal cortex and a negative correlation in multimodal cortex. Furthermore, MEG-FC connectivity strength is negatively correlated in unimodal cortex but positively correlated in multimodal cortex. This complex pattern of correlations may reflect a sophisticated spatiotemporal pattern of brain information processing. During resting-state, external stimuli is minimal, and the brain primarily processes internal information. As a result, rsBEN does not show strong correlations in unimodal cortex, but decreases as the cortical hierarchy increases in multimodal cortex. In the context of watching a movie, sensory information is quickly received and transmitted in unimodal cortex, and then processed in multimodal regions. This results in mvBEN being positively correlated with unimodal cortex and negatively correlated in multimodal cortex. As the positive correlation between SC connection strength and SC-BOLD FC coupling in unimodal cortex may indicate faster information reception and transmission speeds for external stimuli. Information gradually converges in both unimodal and multimodal integration areas, reaching the highest rBEN, and then decreases as it is integrated and internalized.

Some limitations should be noted in this study. The BEN and brain network used were based on group averages rather than individual-level results, which may introduce some potential errors. However, we also analyzed the individual rBEN and functional network connectivity, including connection strength, degree centrality, and local efficiency, for 176 participants, and obtained consistent significant negative correlations. As this is only a draft, further refinements are needed. For example, some clearly U-shaped relationships require quadratic fitting, and rigorous statistical tests need to be conducted for various correlations. Individual-level data should be incorporated, and more comprehensive results and interpretations need to be further organized.

## Acknowledgments

MRI and MEG data were provided by the Human Connectome Project, WU-Minn Consortium (Principal Investigators: David Van Essen and Kamil Ugurbil; 1U54MH091657) funded by the 16 NIH Institutes and Centers that support the NIH Blueprint for Neuroscience Research; and by the McDonnell Center for Systems Neuroscience at Washington University. We thank Golia Shafiei et al for sharing their dataset.

## References

Achard, S. and E. Bullmore (2007). "Efficiency and cost of economical brain functional networks." PLoS computational biology 3(2): e17.

Ajilore, O., M. Lamar and A. Kumar (2014). "Association of brain network efficiency with aging, depression, and cognition." The American Journal of Geriatric Psychiatry 22(2): 102–110.

Bassett, D. S. and O. Sporns (2017). "Network neuroscience." Nature neuroscience 20(3): 353–364.

Bellec, P., V. Perlbarg, S. Jbabdi, M. Pélégrini-Issac, J.-L. Anton, J. Doyon and H. Benali (2006). "Identification of large-scale networks in the brain using fMRI." Neuroimage 29(4): 1231–1243.

Brookes, M. J., J. R. Hale, J. M. Zumer, C. M. Stevenson, S. T. Francis, G. R. Barnes, J. P. Owen, P. G. Morris and S. S. Nagarajan (2011). "Measuring functional connectivity using MEG: methodology and comparison with fcMRI." Neuroimage 56(3): 1082–1104.

Camargo, A., G. Del Mauro and Z. Wang (2024). "Task induced changes in brain entropy." Journal of Neuroscience Research 102(2): e25310.

Catal, Y., A. Wolman, S. Abbasi and G. Northoff (2023). "Intrinsic neural timescales attenuate information transfer along the uni-transmodal hierarchy." bioRxiv: 2023.2007. 2028.551047.

Chang, C. H., S. A. Nastase and U. Hasson (2022). "Information flow across the cortical timescale hierarchy during narrative construction." Proceedings of the National Academy of Sciences 119(51): e2209307119.

Chang, D., D. Song, J. Zhang, Y. Shang, Q. Ge and Z. Wang (2018). "Caffeine caused a widespread increase of resting brain entropy." Scientific reports 8(1): 2700.

Chang, D., J. Zhang, W. Peng, Z. Shen, X. Gao, Y. Du, Q. Ge, D. Song, Y. Shang and Z. Wang (2018). "Smoking cessation with 20 Hz repetitive transcranial magnetic stimulation (rTMS) applied to two brain regions: a pilot study." Frontiers in human neuroscience 12: 344.

Clausius, R. J. E. (1857). Ueber die Art der Bewegung, welche wir Wärme nennen, Verlag nicht ermittelbar.

Del Mauro, G. and Z. Wang (2024). "Associations of brain entropy estimated by resting state fMRI with physiological indices, body mass index, and cognition." Journal of Magnetic Resonance Imaging 59(5): 1697–1707.

Del Mauro, G. and Z. Wang (2024). "rsfMRI-based Brain Entropy is negatively correlated with Gray Matter Volume and Surface Area." bioRxiv: 2024.2004. 2028.591371.

Dong-Hui Song, Y. W., Ze Wang (2024). "Increased Resting Brain Entropy in Mild to Moderate Depression was Decreased by Nonpharmacological Treatment " submitted.

Faust, O., P. C. A. Ang, S. D. Puthankattil and P. K. Joseph (2014). "Depression diagnosis support system based on EEG signal entropies." Journal of mechanics in medicine and biology 14(03): 1450035.

Golesorkhi, M., J. Gomez-Pilar, F. Zilio, N. Berberian, A. Wolff, M. C. Yagoub and G. Northoff (2021). "The brain and its time: intrinsic neural timescales are key for input processing." Communications biology 4(1): 970.

Gómez, C. and R. Hornero (2010). "Entropy and complexity analyses in Alzheimer’s disease: An MEG study." The open biomedical engineering journal 4: 223.

Gruskin, D. C. and G. H. Patel (2022). "Brain connectivity at rest predicts individual differences in normative activity during movie watching." NeuroImage 253: 119100.

Guell, X., J. D. Schmahmann, J. D. Gabrieli and S. S. Ghosh (2018). "Functional gradients of the cerebellum." elife 7: e36652.

Hagmann, P., M. Kurant, X. Gigandet, P. Thiran, V. J. Wedeen, R. Meuli and J.-P. Thiran (2007). "Mapping human whole-brain structural networks with diffusion MRI." PloS one 2(7): e597.

Hansen, J. Y., G. Shafiei, K. Voigt, E. X. Liang, S. M. Cox, M. Leyton, S. D. Jamadar and B. Misic (2023). "Integrating multimodal and multiscale connectivity blueprints of the human cerebral cortex in health and disease." PLoS biology 21(9): e3002314.

Hearne, L. J., J. B. Mattingley and L. Cocchi (2016). "Functional brain networks related to individual differences in human intelligence at rest." Scientific reports 6(1): 1–8.

Herculano-Houzel, S. (2009). "The human brain in numbers: a linearly scaled-up primate brain." Frontiers in human neuroscience 3: 857.

Jiang, W., L. Cai and Z. Wang (2023). "Common hyper-entropy patterns identified in nicotine smoking, marijuana use, and alcohol use based on uni-drug dependence cohorts." Medical & biological engineering & computing 61(12): 3159–3166.

Lake, D. E., J. S. Richman, M. P. Griffin and J. R. Moorman (2002). "Sample entropy analysis of neonatal heart rate variability." American Journal of Physiology-Regulatory, Integrative and Comparative Physiology 283(3): R789–R797.

Langer, N., A. Pedroni, L. R. Gianotti, J. Hänggi, D. Knoch and L. Jäncke (2012). "Functional brain network efficiency predicts intelligence." Human brain mapping 33(6): 1393–1406.

Lau, Z. J., T. Pham, S. A. Chen and D. Makowski (2022). "Brain entropy, fractal dimensions and predictability: A review of complexity measures for EEG in healthy and neuropsychiatric populations." European Journal of Neuroscience 56(7): 5047–5069.

Li, Z., Z. Fang, N. Hager, H. Rao and Z. Wang (2016). "Hyper-resting brain entropy within chronic smokers and its moderation by Sex." Scientific reports 6(1): 29435.

Lin, C., S.-H. Lee, C.-M. Huang, G.-Y. Chen, P.-S. Ho, H.-L. Liu, Y.-L. Chen, T. M.-C. Lee and S.-C. Wu (2019). "Increased brain entropy of resting-state fMRI mediates the relationship between depression severity and mental health-related quality of life in late-life depressed elderly." Journal of affective disorders 250: 270–277.

Lin, L., D. Chang, D. Song, Y. Li and Z. Wang (2022). "Lower resting brain entropy is associated with stronger task activation and deactivation." NeuroImage 249: 118875.

Liu, P.-S., D.-H. Song, X.-P. Deng, Y.-Q. Shang, G. Qiu, Z. Wang and H. Zhang (2024). "Intermittent theta burst stimulation (iTBS)-induced changes of resting-state brain entropy (BEN)." bioRxiv: 2024.2005. 2015.591015.

Liu, X., D. Song, Y. Yin, C. Xie, H. Zhang, H. Zhang, Z. Zhang, Z. Wang and Y. Yuan (2020). "Altered Brain Entropy as a predictor of antidepressant response in major depressive disorder." Journal of Affective Disorders 260: 716–721.

Liu, Z.-Q., G. Shafiei, S. Baillet and B. Misic (2023). "Spatially heterogeneous structure-function coupling in haemodynamic and electromagnetic brain networks." NeuroImage 278: 120276.

Luo, L. (2021). "Architectures of neuronal circuits." Science 373(6559): eabg7285.

Margulies, D. S., S. S. Ghosh, A. Goulas, M. Falkiewicz, J. M. Huntenburg, G. Langs, G. Bezgin, S. B. Eickhoff, F. X. Castellanos and M. Petrides (2016). "Situating the default-mode network along a principal gradient of macroscale cortical organization." Proceedings of the National Academy of Sciences 113(44): 12574–12579.

Menon, V. (2011). "Large-scale brain networks and psychopathology: a unifying triple network model." Trends in cognitive sciences 15(10): 483–506.

Park, H.-J. and K. Friston (2013). "Structural and functional brain networks: from connections to cognition." Science 342(6158): 1238411.

Pincus, S. M. (1991). "Approximate entropy as a measure of system complexity." Proceedings of the national academy of sciences 88(6): 2297–2301.

Pincus, S. M. (2006). "Approximate entropy as a measure of irregularity for psychiatric serial metrics." Bipolar disorders 8(5p1): 430-440.

Power, J. D., A. L. Cohen, S. M. Nelson, G. S. Wig, K. A. Barnes, J. A. Church, A. C. Vogel, T. O. Laumann, F. M. Miezin and B. L. Schlaggar (2011). "Functional network organization of the human brain." Neuron 72(4): 665–678.

Richman, J. S. and J. R. Moorman (2000). "Physiological time-series analysis using approximate entropy and sample entropy." American journal of physiology-heart and circulatory physiology 278(6): H2039–H2049.

Sabeti, M., S. Katebi and R. Boostani (2009). "Entropy and complexity measures for EEG signal classification of schizophrenic and control participants." Artificial intelligence in medicine 47(3): 263–274.

Schaefer, A., R. Kong, E. M. Gordon, T. O. Laumann, X.-N. Zuo, A. J. Holmes, S. B. Eickhoff and B. T. Yeo (2018). "Local-global parcellation of the human cerebral cortex from intrinsic functional connectivity MRI." Cerebral cortex 28(9): 3095–3114.

Schröter, M., O. Paulsen and E. T. Bullmore (2017). "Micro-connectomics: probing the organization of neuronal networks at the cellular scale." Nature Reviews Neuroscience 18(3): 131–146.

Senkowski, D. and A. K. Engel (2024). "Multi-timescale neural dynamics for multisensory integration." Nature Reviews Neuroscience 25(9): 625–642.

Shafiei, G., S. Baillet and B. Misic (2022). "Human electromagnetic and haemodynamic networks systematically converge in unimodal cortex and diverge in transmodal cortex." PLoS biology 20(8): e3001735.

Shannon, C. E. (1948). "A mathematical theory of communication." The Bell system technical journal 27(3): 379–423.

Song, D.-H., X.-P. Deng, Y.-Q. Shang, D. Chang and Z. Wang (2024). "Altered resting-state brain entropy (BEN) by rTMS across the human cortex." bioRxiv: 2024.2007. 2016.601109.

Song, D.-H. and Z. Wang (2024). "Regional Brain Entropy During Movie-watching." bioRxiv: 2024.2006. 2012.598767.

Song, D.-H. and Z. Wang (2024). "The Relationships of Resting-state Brain Entropy (BEN), Ovarian Hormones and Behavioral Inhibition and Activation Systems (BIS/BAS)." bioRxiv: 2024.2006. 2004.595915.

Song, D., D. Chang, J. Zhang, Q. Ge, Y.-F. Zang and Z. Wang (2019). "Associations of brain entropy (BEN) to cerebral blood flow and fractional amplitude of low-frequency fluctuations in the resting brain." Brain imaging and behavior 13(5): 1486–1495.

Song, D., D. Chang, J. Zhang, W. Peng, Y. Shang, X. Gao and Z. Wang (2019). "Reduced brain entropy by repetitive transcranial magnetic stimulation on the left dorsolateral prefrontal cortex in healthy young adults." Brain imaging and behavior 13: 421–429.

Song, D. and Z. Wang (2024). "Neurotransmitters Contribute Structure-Function Coupling: Evidence from Grey Matter Volume (GMV) and Brain Entropy (BEN)." bioRxiv: 2024.2009. 2007.611832.

Song, D. and Z. Wang (2024). "Phenotype Prediction Using BEN-Based Predictive Modeling (BPM)." bioRxiv: 2024.2009. 2007.611838.

Song Donghui, C. D., Wang Ze (2023). "Reproducible brain entropy increases during ruminations." The Organization for Human Brain Mapping (OHBM) 2023 Annual Meeting.

Song, M., Y. Zhou, J. Li, Y. Liu, L. Tian, C. Yu and T. Jiang (2008). "Brain spontaneous functional connectivity and intelligence." Neuroimage 41(3): 1168–1176.

Stam, C. J. (2005). "Nonlinear dynamical analysis of EEG and MEG: review of an emerging field." Clinical neurophysiology 116(10): 2266–2301.

Suárez, L. E., R. D. Markello, R. F. Betzel and B. Misic (2020). "Linking structure and function in macroscale brain networks." Trends in cognitive sciences 24(4): 302–315.

Tian, Y., D. S. Margulies, M. Breakspear and A. Zalesky (2020). "Topographic organization of the human subcortex unveiled with functional connectivity gradients." Nature neuroscience 23(11): 1421–1432.

Van Den Heuvel, M. P., C. J. Stam, R. S. Kahn and H. E. H. Pol (2009). "Efficiency of functional brain networks and intellectual performance." Journal of Neuroscience 29(23): 7619–7624.

Van Essen, D. C., S. M. Smith, D. M. Barch, T. E. Behrens, E. Yacoub, K. Ugurbil and W.-M. H. Consortium (2013). "The WU-Minn human connectome project: an overview." Neuroimage 80: 62–79.

Vázquez-Rodríguez, B., L. E. Suárez, R. D. Markello, G. Shafiei, C. Paquola, P. Hagmann, M. O. Van Den Heuvel, B. C. Bernhardt, R. N. Spreng and B. Misic (2019). "Gradients of structure–function tethering across neocortex." Proceedings of the National Academy of Sciences 116(42): 21219–21227.

Von Bartheld, C. S., J. Bahney and S. Herculano Houzel (2016). "The search for true numbers of neurons and glial cells in the human brain: A review of 150 years of cell counting." Journal of Comparative Neurology 524(18): 3865–3895.

Vos de Wael, R., O. Benkarim, C. Paquola, S. Lariviere, J. Royer, S. Tavakol, T. Xu, S.-J. Hong, G. Langs and S. Valk (2020). "BrainSpace: a toolbox for the analysis of macroscale gradients in neuroimaging and connectomics datasets." Communications biology 3(1): 103.

Wang, X.-J. (2020). "Macroscopic gradients of synaptic excitation and inhibition in the neocortex." Nature reviews neuroscience 21(3): 169–178.

Wang, Z. (2021). "The neurocognitive correlates of brain entropy estimated by resting state fMRI." NeuroImage 232: 117893.

Wang, Z. (2021). "Resting state fMRI-based temporal coherence mapping." arXiv preprint arXiv:2109.00146.

Wang, Z. and A. s. D. N. Initiative (2020). "Brain entropy mapping in healthy aging and Alzheimer’s disease." Frontiers in Aging Neuroscience 12: 596122.

Wang, Z., Y. Li, A. R. Childress and J. A. Detre (2014). "Brain entropy mapping using fMRI." PloS one 9(3): e89948.

Wolff, A., N. Berberian, M. Golesorkhi, J. Gomez-Pilar, F. Zilio and G. Northoff (2022). "Intrinsic neural timescales: temporal integration and segregation." Trends in cognitive sciences 26(2): 159–173.

Xue, S.-W., Q. Yu, Y. Guo, D. Song and Z. Wang (2019). "Resting-state brain entropy in schizophrenia." Comprehensive Psychiatry 89: 16–21.

Yao, Y., W. Lu, B. Xu, C. Li, C. Lin, D. Waxman and J. Feng (2013). "The increase of the functional entropy of the human brain with age." Scientific reports 3(1): 2853.

Yeh, C. H., D. K. Jones, X. Liang, M. Descoteaux and A. Connelly (2021). "Mapping structural connectivity using diffusion MRI: challenges and opportunities." Journal of Magnetic Resonance Imaging 53(6): 1666–1682.

Zhao, Z., Y. Shuai, Y. Wu, X. Xu, M. Li and D. Wu (2024). "Age-dependent functional development pattern in neonatal brain: an fMRI-based brain entropy study." NeuroImage: 120669.

